# Inosine Triphosphate Pyrophosphatase as a Candidate Associated with White Matter Abnormalities in Chronic Heart Failure

**DOI:** 10.64898/2026.11.03.764123

**Authors:** Kouki Takeuchi, Hideaki Suzuki, Akira Sumiyoshi, Ryo Kurosawa, Ai Sugano, Takuya Urushihata, Marina Arai, Kotaro Nochioka, Kaoru Ito, Akiko Satoh, Toru Nakazawa, Satoshi Yasuda

## Abstract

**Background:** White matter hyperintensities (WMHs) are common with normal brain aging and frequently observed also in chronic heart failure (CHF). However, the relationship between CHF and white matter abnormalities and the biological factors remain unclear. We investigated WMH and associated white matter microstructural changes, such as fractional anisotropy (FA), in CHF and explored candidate molecular factors.

**Methods:** We conducted a longitudinal prospective observation including brain T2-FLAIR magnetic resonace imaging for 32 asymptomatic Stage B (mean age 63.3±10.5 years; 25% women) and 23 symptomatic Stage C CHF patients (mean age 67.3±7.1 years; 30% women). Plasma proteomics and Mendelian randomization (MR) were used to identify candidate proteins associated with both CHF and WMH. The association of candidate proteins was further investigated using complementary analyses in CHF rats, cultured cortical neurons, and community-based genetic data in UK Biobank.

**Results:** Global WMH volume increased more in Stage C than Stage B patients, predominantly near the anterior periventricular white matter, despite comparable cardiac indices (4,717±5,335 mm³ vs. 2,116±3,134 mm³, P=0.027). SomaScan-based proteomic analysis of plasma samples from this patient cohort revealed that 44 proteins are significantly associated with both Stage C CHF and WMH progression (P<0.05). Of these 44 proteins, two-step Mendelian randomization analysis identified inosine triphosphate pyrophosphatase (ITPA) as the only candidate protein, further supported by an association between rare variants in ITPA and FA. In CHF rats, the anterior pericallosal cortex showed decreases in myelin basic protein density and FA and an increase in ITPA expression compared to sham rats. In cultured cortical neurons, recombinant ITPA exposure reduced total axonal length.

**Conclusions:** Symptomatic CHF was associated with WMH and impaired white matter integrity. ITPA emerged as a candidate factor across human, animal, and cellular analyses. These findings provide a basis for investgating the biological pathways linking CHF and white matter injury.

## Introduction

Chronic heart failure (CHF), characterized by systemic hypoperfusion and neurohumoral activation,^1^ is closely associated with cerebral dysfunction, including dementia^2^ and depression.^3^ Cerebral hypoperfusion and possible resultant damage have been proposed as potential mechanisms underlying brain structural abnormalities and neuropsychiatric disorders.^4–6^ These observations suggest that chronic cardiovascular dysfunction may contribute to structural brain injury, although the mechanisms linking CHF to cerebral abnormalities remain incompletely understood.

White matter hyperintensities (WMHs), detected as hyperintense lesions on T2-weighted magnetic resonance imaging (MRI), are common manifestations of cerebral small vessel disease affecting the cerebral microvasculature and can lead to ischemic and hemorrhagic stroke, dementia, and even death.^7–9^ WMHs may reflect various pathological changes, such as loss of myelin, microinfarcts, and microhemorrhages.^10^ WMH has classically been attributed to cerebral small vessel damage and subsequent hypoperfusion associated with aging and atherosclerosis.^11,12^ Several mechanisms, including neurovascular inflammation^13^ and genetic variants,^14^ have been implicated in WMH progression. However, the biological underpinning of WMH is poorly understood and mechanism-based treatments are not available currently.

Previous cross-sectional studies have reported inconsistent associations between CHF and WMH,^15–17^ and longitudinal evidence is limited. Moreover, the molecular factors that may link CHF to white matter abnormalities have not been well characterized. We therefore examined longitudinal changes in WMH and white matter integrity, such as fractional anisotropy (FA), across different stages of CHF using neuroimaging. White matter FA may be a more sensitive imaging marker of subtle white matter injury beyond the abnormalities visible as WMH on conventional T2-weighted MRI.^18^ We hypothesized that more severe heart failure would exhibit greater progression of white matter abnormalities. In addition, histological analyses in an animal model of CHF were conducted for complementary analysis. We further explored candidate proteins related to CHF-associated WMH progression through plasma proteomic screening and Mendelian randomization (MR) analysis for causal inference, followed by complementary animal, cell culture and community-based cohort data to investigate their potential relevance to white mater abnormalities.

## Methods

### Study Population

The clinical study protocol was approved by the Tohoku University Ethics Committee (2025-1-835) and was registered in the University Hospital Medical Information Network as Brain Assessment and Investigation in Heart Failure Trial 2 (B-HeFT2), a longitudinal study of brain structure and function in patients stratified by heart failure severity (UMIN000020355). All procedures of animal study were approved by the Animal Care Committee at Tohoku University (2023AcA-007-02; 2025MdA-033). Imaging representation of community-based cohort (UK Biobank) was conducted under terms of UKB access approval 18545 and 94113.

### CHF Cohort

The study cohort was derived from a prospective observational study of patients with CHF at Tohoku university hospital.^6^ Baseline brain MRI, cine cardiovascular magnetic resonance (CMR), echocardiography and blood sampling were initially performed in 40 patients with asymptomatic Stage B and 40 patients with symptomatic Stage C CHF, aged 45–90 years, from August 2012 to April 2013, based on the 2009 American College of Cardiology/American Heart Association guidelines for the diagnosis and management of heart failure.^19^ The sample size was determined by the availability of paired baseline and follow-up brain MRI data from our cohort. Of these 80 patients, 25 were excluded from the follow-up analysis: 8 patients with Stage B CHF (7 were lost to follow-up and 1 died) and 17 patients with Stage C CHF (9 were lost to follow-up, 6 died, and 2 underwent cardiac device implantation). Consequently, 55 patients (32 with Stage B HF and 23 with Stage C HF) underwent follow-up brain MRI after a median follow-up period of 3.1 years. Serial brain MRI included 3D-T2 fluid-attenuated inversion-recovery (FLAIR) images for assessment of WMH. Follow-up plasma samples were also collected from the 55 patients, and plasma samples obtained at both baseline and follow-up were used for proteomic analysis.

### Brain and cardiac magnetic resonance imaging acquisition and pre-processing

Brain MRI was performed using a 1.5-T whole-body MRI system (Signa HDxt, GE Medical Systems, Milwaukee, WI, USA) and an 8-channel NVArray coil (coil selection using head-A). Cube sagittal 3D-T2 FLAIR images were acquired for WMH measurements with the following parameters: fast-recovery fast spin-echo with variable flip angle refocusing; repetition time 8000msec; echo time 117.9msec, inversion time 2253msec; echo train length of 140; field of view 220mm; acquisition matrix of 224×224; 192 sections; voxel size 1.0×0.98×0.98mm. WMH was manually segmented on 3D-T2 FLAIR images, and global WMH volumes were quantified. For voxel-wise analysis of spatial changes in WMH, WMH masks were co-registered to T1-weighted images, spatially normalized to Montreal Neurological Institute space, and longitudinal WMH change maps were generated.

Left ventricular cardiac and stroke volume indices were derived from cine cardiovascular magnetic resonance imaging. The same MRI scanner, acquisition protocol, and image-processing procedures were used for our cohort to ensure comparability. WMH segmentation and volume quantification were performed using the same assessment procedures for all images (see **Supplementary material online, Method S1** and **S2**).

### Proteomics measurements and Protein-wise association analyses

Plasma samples collected from participants in B-HeFT2 were analyzed using the SomaScan Discovery Platform V4.1 (SomaLogic Inc., Boulder, CO, USA), an established aptamer-based platform for large-scale proteomic analyses in cardiovascular disease research.^20^ SomaScan is based on slow off-rate modified single-stranded DNA aptamers (SOMAmers). SOMAmers are designed to bind to proteins based on a particular sequence or three-dimensional structure, thereby exhibiting high specificity and affinity for proteins and minimize nonspecific binding interactions owing to their slow dissociation rates. Additional details on the assay and its performance have been previously published.^21^

All plasma samples were analyzed using the same version of the SomaScan assay and processed in a single batch to ensure consistency and to minimize potential batch effects. The SomaScan assay was conducted according to the standardized quality control processes of SomaLogic Inc., with 7,596 SOMAmer reagents measured. A total of 307 control or non-human-targeting reagents were not included in our analysis, and an additional 158 reagents were excluded based on the manufacturer’s quality control flags, leaving 7,131 proteins in subsequent analyses.

Protein-wise association analyses were performed separately for WMH progression and Stage C CHF. For each of the 7,131 proteins, plasma protein levels were log-transformed before analysis. For WMH progression, changes in protein levels from baseline to follow-up were calculated and examined for their association with changes in global WMH volume between the same two time points using age– and sex-adjusted linear regression models. For Stage C CHF, changes in protein levels from baseline to follow-up were compared between patients with Stage B and Stage C CHF using age– and sex-adjusted logistic regression models. Proteins showing evidence of association with both phenotypes (P<0.05) were prioritized as candidate proteins for MR analyses subsequently.

### Two-step Mendelian randomization analysis

We applied a sequential two-step MR framework to assess whether candidate proteins lie on the putative causal pathway linking CHF and WMH burden. The first MR estimated the causal effect of CHF on circulating protein levels (β₁), while the second estimated the causal effect of circulating protein levels on WMH burden (β₂). To avoid bias arising from sample overlap, two-sample MR was performed using independent genome-wide association study (GWAS) summary statistics. Summary statistics for CHF were obtained from FinnGen Freeze 10,^22^ WMH from a GWAS meta-analysis of WMHs of the UK Biobank and CHARGE consortium cohorts,^13^ and plasma protein levels from the deCODE proteomic GWAS.^23^ Independent instrumental variables were selected by linkage disequilibrium clumping (P<5×10⁻⁸, r²<0.05, 10,000-kb window, minor allele frequency ≥0.01). The Inverse-Variance Weighted (IVW) method was used as the primary MR analysis, with the weighted median, simple mode, weighted mode, and MR-Egger regression performed as sensitivity analyses. The MR-Egger intercept test was used to assess directional horizontal pleiotropy.

### Rat model and tissue preparation

A myocardial infarction model was used as a rat model of CHF. Animals were allocated to ex vivo MRI and histological analyses (14 CHF and 14 sham rats) or Western blot analysis (5 CHF and 5 sham rats). For ex vivo MRI and histological analyses, rats were deeply anesthetized with 5% isoflurane and perfused via the left ventricle, following incision of right atrium, with 100mL of phosphate-buffered saline (PBS), 0.1mol/l, pH 7.4, containing heparin at a final concentration of 50U/mL prepared from a 1,000USP U/mL stock solution. Perfusion of 200mL of 4% paraformaldehyde (PFA) with 2mM prohance was followed by that of PBS.^4^ After perfusion, the rats were decapitated, and the skin, lower jaw, ears, and cartilaginous portion of the nose were removed while the brain was retained within the skull. The specimens were post-fixed overnight at 4°C in 4% PFA containing 2 mM prohance and subsequently stored in PBS containing 2 mM prohance. For western blotting analysis, rats were deeply anesthetized with 5% isoflurane and perfused with PBS containing heparin alone, followed by decapitated.

### Brain magnetic resonance imaging recordings in animals

Ex vivo MRI was performed at QST using a 7-T BioSpec MRI system (Bruker BioSpin MRI GmbH, Ettlingen, Germany) operated with ParaVision 7.0.0. Each skull-containing specimen was placed in a 30-mL syringe, which was sealed to prevent desiccation. Radiofrequency transmission was performed using an actively detunable volume transmit coil with an inner diameter of 86mm (T12053V3; Bruker BioSpin), and signal reception was performed using an eight-channel receive-only array coil with an inner diameter of 37mm (RAPID Biomedical GmbH, Rimpar, Germany). As spontaneous white matter lesions resembling age-related WMH in humans are uncommon in rats during their normal lifespan,^24^ diffusion tensor imagings (DTIs) were acquired for the assessment of white matter injuries using the Bruker three-dimensional spin-echo diffusion sequence (DtiStandard) with the following parameters: repetition time/echo time, 100/22ms; field of view, 30×18×22 mm³; acquisition matrix, 150×90×110; isotropic spatial resolution, 200µm; 30 diffusion-encoding directions at a b-value of 800s/mm²; one non-diffusion-weighted image (b=0s/mm²); diffusion gradient duration (δ), 4.5 ms; diffusion gradient separation (Δ), 10.6ms; four signal averages; and total acquisition time, 25h 45min 52s. Bruker raw data were imported into DSI Studio and converted to NIfTI format together with the corresponding b-values and b-vectors. Diffusion-weighted images were subsequently corrected for eddy-current-induced distortions using FSL’s eddy_correct, with the first volume as the reference and spline interpolation. Diffusion tensors were fitted using FSL’s dtifit to generate fractional anisotropy (FA) maps, which were used for region of interest (ROI) analyses.

### Brain magnetic resonance imaging analysis in animals

FA values within the ROIs were calculated based on previously published methods.^25^ The ROIs were selected within the anterior pericallosal region, which corresponded to the approximate anatomical area identified in the voxel-wise analysis of CHF cohort. In rodents, unlike in primates, the pericallosal region adjacent to the corpus callosum contains a prominent myelin-rich stratum and is considered anatomically comparable to the corresponding white matter region in humans.^26,27^ Therefore, the pericallosal region was used as a representative white matter region for the assessment of white matter injury in the present rodent model. Bilateral ROIs were manually delineated on the basis of FA maps using the ITK-SNAP version 4.0.2.^28^ Each ROI was drawn on five sequential slices (1000μm) of the rigidly aligned FA maps to generate the complete ROI. The mean FA value within each ROI was obtained using the segmentation statistics function in ITK-SNAP.

### Analysis of Community-based cohort

Publicly available gene-based association results were queried using GeneBass, an online resource containing exome-based association statistics for 4,529 phenotypes derived from up to 394,841 UK Biobank participants.^29^ The sequence kernel association test–optimal (SKAT-O) for predicted loss-of-function variants in ITPA and weighted-mean FA across 27 white matter tracts were examined to measure the P values, as previously described.^30^ The corresponding burden-test effect estimate was recorded to indicate the direction and magnitude of the association. Burden tests are powerful when most variants in a region are causal and the effects are in the same direction, whereas SKAT-O adaptively combines the burden and the sequence kernel association tests, thereby maintaining power when qualifying variants have either largely concordant effects or heterogeneous effects in magnitude and direction.^31^ The phenotype-specific analysis included 16,496 participants with available data. The P value calculated by SKAT-O displayed in GeneBass was recorded. GeneBass was accessed on July 15, 2026.

For illustrative purposes, probabilistically defined trajectories of the 27 white matter tracts shown in **Figure 4** were generated as previously described.^32^ Briefly, surface meshes were made from probabilistic descriptions of 27 white matter tracts in one UK Biobank participant. The tracts meshes were rendered as a 3D fashion onto a transparent brain mask that was derived from the MRI template available in SPM 12. Tracts were classified as association fibers, thalamic radiations, and projection fibers according to a previously published classification.^33^

### Statistical Analysis

Continuous variables were expressed as mean ± standard deviation. Normality was tested using the Shapiro-Wilk test (α=0.05). When the data followed a normal distribution, Student’s t-test was used as appropriate. For non-normally distributed data, Mann–Whitney U test was applied. Categorical variables were expressed as percentages using with Fisher’s exact test or the χ^2^ test, as appropriate. Associations between two continuous variables were evaluated using Pearson’s correlation analysis. Statistical analyses were performed using Stata statistical software version 18 (StataCorp) with a two-tailed significance threshold of P<0.05 unless otherwise stated.

We first examined whether WMH changes differed between the Stage B and Stage C groups in CHF cohort to investigate the association between CHF and global changes in WMH. Differences in WMH changes were calculated by subtracting the baseline value from the follow-up value using global volumetry. Whole-brain voxel-wise analysis in which each voxel of the normalized white matter map was also used as the dependent variable with CHF stage as the primary explanatory variable and adjustment for age and sex. A statistical significance for the voxel-wise analysis was defined at *P*<0.05 after family-wise error (FWE) correction for multiple comparisons.^34^

We next tested a combination of plasma proteomics in the 55 CHF patients and MR analysis in external cohorts to explore candidate proteins associated both with CHF and WMH progression. No correction for multiple testing was applied to either the proteomic or MR analyses; the significance thresholds were considered exploratory. Among these, inosine triphosphate pyrophosphatase (ITPA) was further investigated for experimental analyses. The association between ITPA and white matter injury was also examined using the GeneBass rare variant analysis, given the marginal statistical significance observed in the IVW MR analysis. Loss-of-function burden of ITPA was predicted for FA in 27 white matter tracts at a significance threshold of *P*<0.05 after Bonferroni correction for 27 multiple comparisons, corresponding to a raw P value of <0.00185 (n=16496).

We finally performed additional examinations using animals to corroborate the relationships among CHF, WMH progression, and ITPA. To investigate the relationship between CHF and WMH, myelin basic protein (MBP) immunodensity obtained from immunohistochemistry and FA in the pericallosal cortex were compared between CHF and sham rats. To examine ITPA expression in CHF, cerebral ITPA protein levels were compared between CHF and sham rats using Western blot analysis. To assess the effect of ITPA on axonal morphology, total axonal length was compared between cells treated with recombinant ITPA and those treated with vehicle in the cultured neuron experiment.

The observational component of this study was reported in accordance with the Strengthening the Reporting of Observational Studies in Epidemiology (STROBE) guidelines, ^35^ whereas the Mendelian randomization analysis was reported in accordance with the STROBE-MR guidelines.^36^

## Results

### WMH progression in patients with Stage B and Stage C CHF

Baseline characteristics in Stage B and Stage C groups were shown in **Table 1**. Baseline demographic characteristics, vascular risk factors, total WMH volume, and intracranial volume were comparable between the two groups (P>0.05). Echocardiography and CMR showed that left ventricular ejection fraction (LVEF) was lower and left ventricular diameter and volume were larger in Stage C group compared to Stage B group (P<0.05). In contrast, left ventricular cardiac indices and systolic volume indices did not differ significantly between the groups (P>0.05).

**Table 1.**
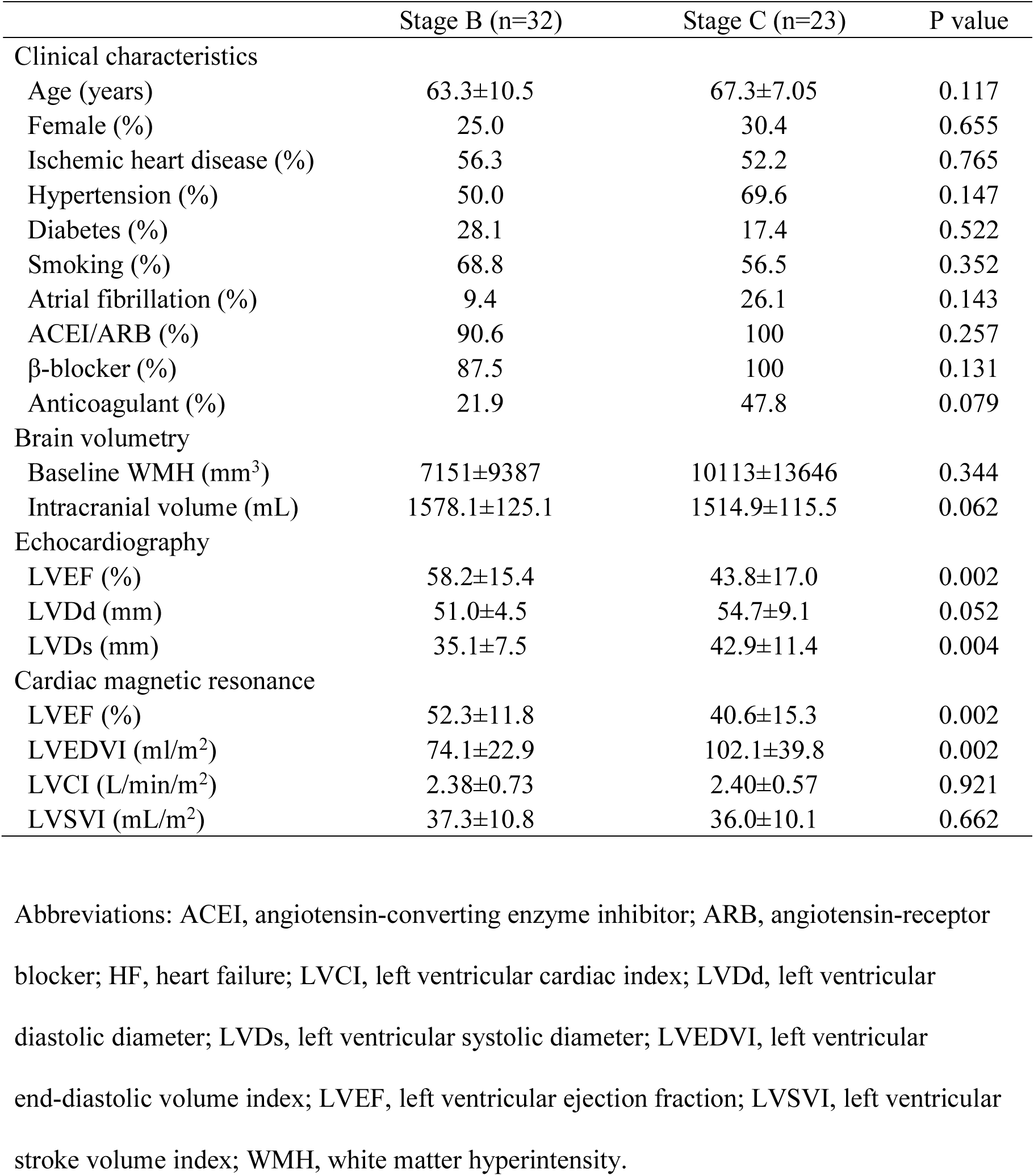
Baseline characteristics.

During a median follow-up of 3.1 years, the increase in global WMH volume was greater in Stage C than in Stage B groups (4717.3±5334.8 vs. 2116.3±3134.4mm^3^, P=0.027, **Figure 1A**). The spatial distribution of WMH increase also differed between the groups, with more extensive changes observed in the Stage C (**Figure 1C**, red regions) compared to Stage B groups (**Figure 1B**, blue regions). In a whole-brain voxel-wise analysis, WMH increase differed significantly between Stage B and Stage C groups near the anterior periventricular white matter (*P*<0.05 [FWE corrected], **Figure 1C**, yellow regions and arrowheads). The change in global WMH volume was not significantly associated with left ventricular cardiac indices or systolic volume indices (P>0.05, see **Supplementary data online, Table S1**).

**Figure 1.**
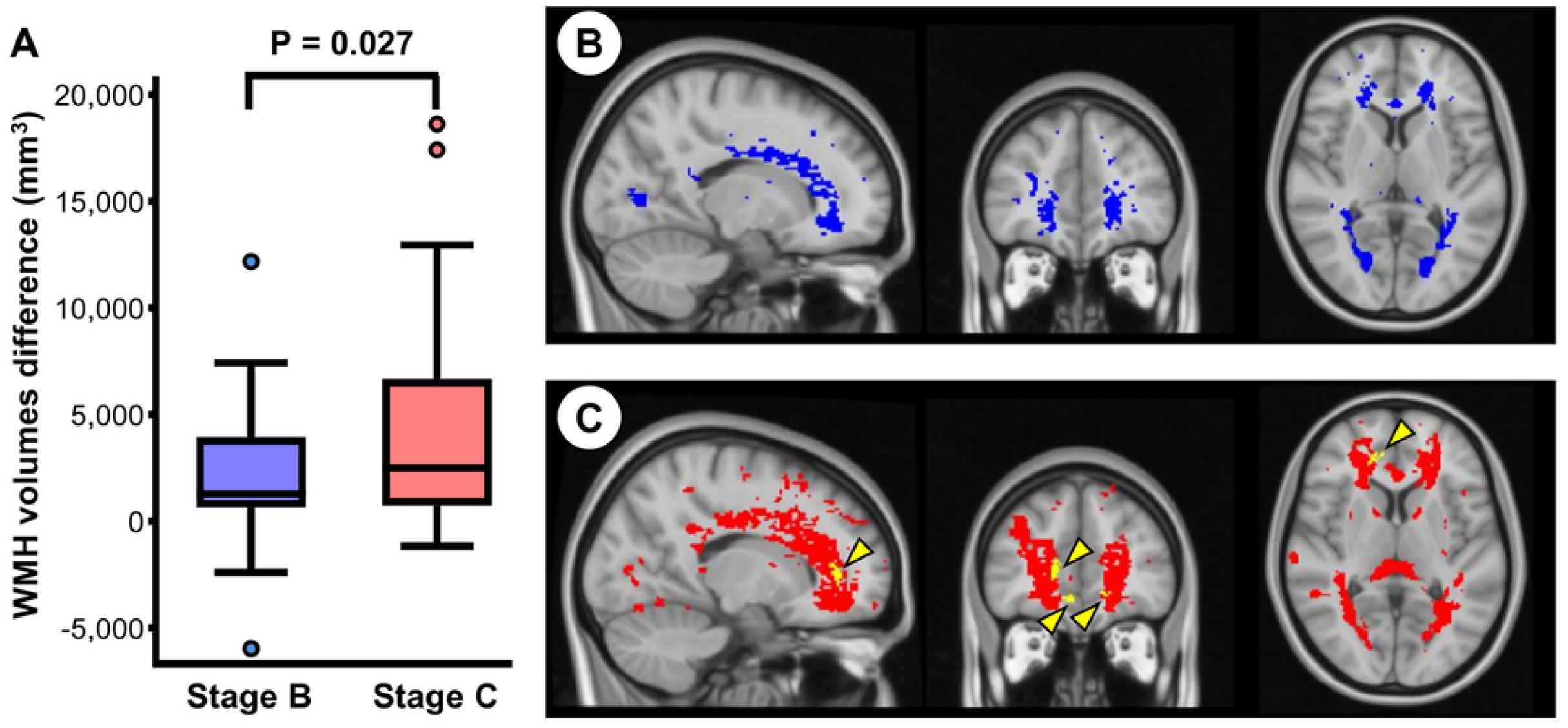
Differences in white matter hyperintensity (WMH) volume change between Stage B and C groups in B-HeFT2. Changes in global WMH volume in Stage B and C groups (n=32 and 23, respectively). Dots represent individual data. Statistical analysis was performed using Student’s t-test (A). Regions with the WMH increase in Stage B (blue regions, B) and C groups (red regions, C). Yellow regions and arrowheads indicate the WMH volume changes near the anterior periventricular white matter, where there was a significant difference between Stage B and C groups in the WMH increase (*P*<0.05, with family-wise error corrections for multiple comparisons).

### Differences in the anterior pericallosal myelination between CHF and sham rats

To complement the longitudinal differences in WMH volume near the anterior periventricular white matter in CHF patients, we performed imaging and histological analyses of the pericallosal myelination in the anterior brain regions in the rat model of CHF (**Figure 2A and C)**. Consistent with the clinical CHF cohort, left ventricular diastolic and systolic diameters were larger and fractional shortening was lower in CHF compared to sham rats (P<0.05, see **Supplementary data online, Table S2**). The analysis of brain DTI images showed that FA in the anterior pericallosal regions, which corresponded to the approximate anatomical area identified in the voxel-wise analysis of CHF cohort, was lower in CHF compared to sham rats (0.297±0.012 vs. 0.320±0.021, P=0.002, **Figure 2B**). Consistently, immunohistochemical analysis also showed that immune density of anterior pericallosal MBP was lower in CHF compared to sham rats (0.377±0.460 vs. 1.000±0.625, P=0.021, **Figure 2D**). There was a significant positive correlation between FA and MBP immunodensity in the corresponding anterior pericallosal regions (R=0.485, P=0.009, **Figure 2E**). Together, these observations support the presence of CHF-associated microstructural and myelin-related abnormalities, which were consistent findings with WMH changes in patients with CHF, in animals.

**Figure 2.**
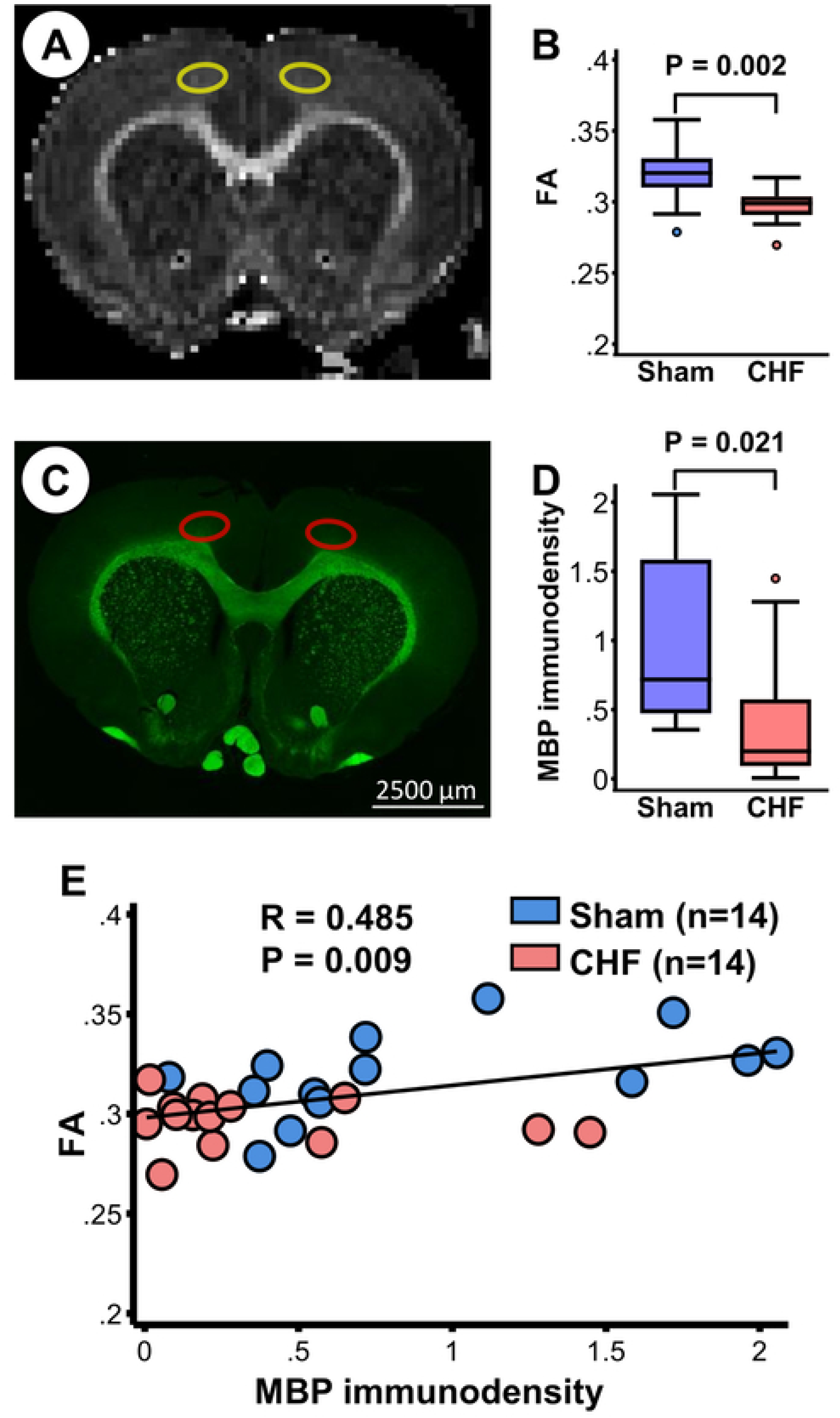
White matter damage induced by chronic heart failure (CHF). A representative image of brain coronal section 1.3mm anterior to bregma by fractional anisotropy (FA) maps in cortex (yellow) (A). Quantification of FA in cortex of sham and CHF rats (n=14, each) (B). A representative image of brain coronal section 1.3mm anterior to bregma with myelin binding protein (MBP) stain in cortex (red), in a region corresponding to the approximate anatomical area identified in the CHF cohort. Scale bar; 2500μm (C). Quantification of MBP immunodensity in cortex of sham and CHF rats normalized to each sham animals processed on the same experimental day (n=14, each) (D). Correlation between MBP immunodensity and FA (E). Statistical analysis was performed using Mann–Whitney U test (B), Student’s t-test (D), and Pearson’s correlation analysis (E).

### Identification of ITPA as a candidate protein associated with both CHF and WMH

To screen for proteins associated with the increased WMH volume in CHF cohort, we performed proteomic analysis to test whether 7,131 proteins were associated both with Stage C CHF and longitudinal WMH progression. Of these, 44 proteins were associated both with the two phenotypes in age– and sex-adjusted analyses (P<0.05, **Figure 3A**). Subsequently, two-step MR was performed for these 44 proteins to investigate their potential causal relationships, assessing CHF-to-protein effects in step 1 and protein-to-WMH effects in step 2. Among the 44 proteins, only ITPA showed a significant causal effect in step 1 and a trend toward significance in step 2 (IVW β_1_=0.098, P=0.025. β_2_=0.009, P=0.092, **Figure 3B**). These findings suggest a potential pathway from the CHF phenotype to increased ITPA levels and that from increased ITPA levels to WMH burden. Although the IVW significance of the association between ITPA levels and WMH burden was marginal, MR-Egger analysis further suggested a potential causal effect of ITPA on WMH progression (β=0.015, P=0.038, see **Supplementary data online, figure S1** and **table S4**). The GeneBass rare variant analysis also showed the ITPA-related white matter abnormality: ITPA-predicted loss-of-function burden was associated with weighted-mean FA in the left superior longitudinal fasciculus (corrected *P*=0.022 [raw P<0.001], **Figure 4**). An association with marginal significance was observed also in the contralateral right superior longitudinal fasciculos (corrected *P*=0.076 [raw P<0.001]). Their effect estimates (beta) calculated by burden test were positive (β=0.0071 and 0.0032), although not statistically significant (corrected *P*>0.05, see **Supplementary data online, Table S5**). These findings suggest that loss of ITPA function may exert a protective effect against white matter abnormalities, as indicated by higher FA values. Given the exploratory nature of the proteomic and MR analyses, the relationships among CHF, WMH progression, and ITPA were further investigated using subsequent complementary experiments.

**Figure 3.**
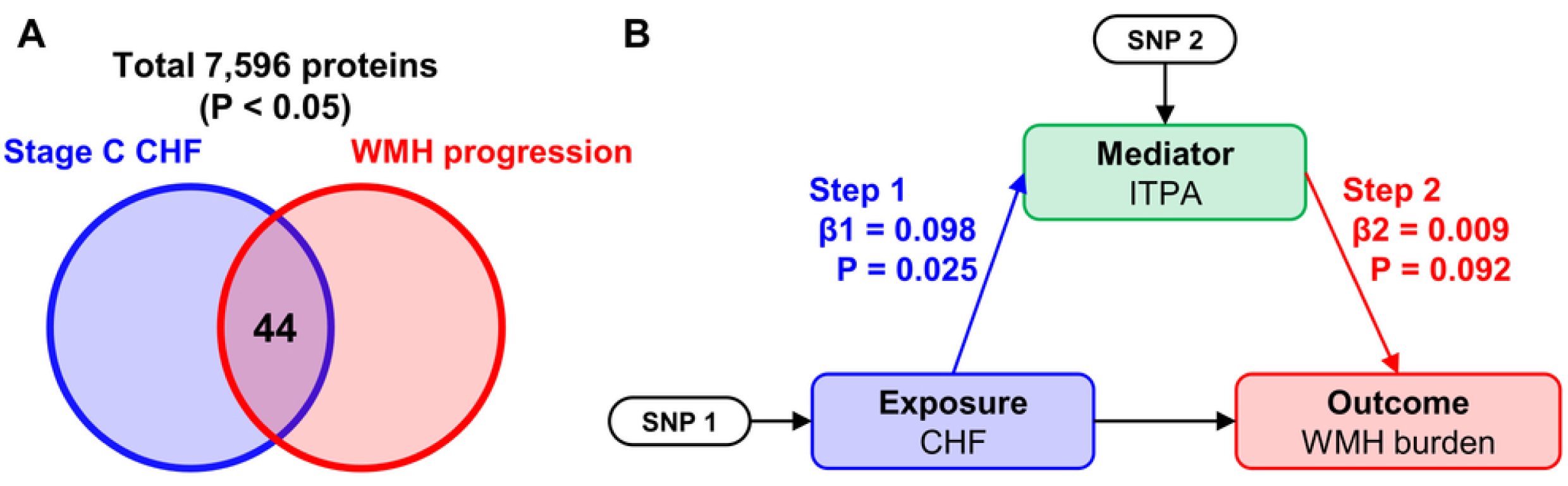
Schematic representation of proteomic analysis and Mendelian randomization. Venn diagram showing overlaps among proteins associated with stage C CHF and WMH progression (A). Results of two-step Mendelian randomization study for inosine triphosphate pyrophosphatase (B). β denotes the inverse-variance weighted method effect estimate for each step of the two-step.

**Figure 4.**
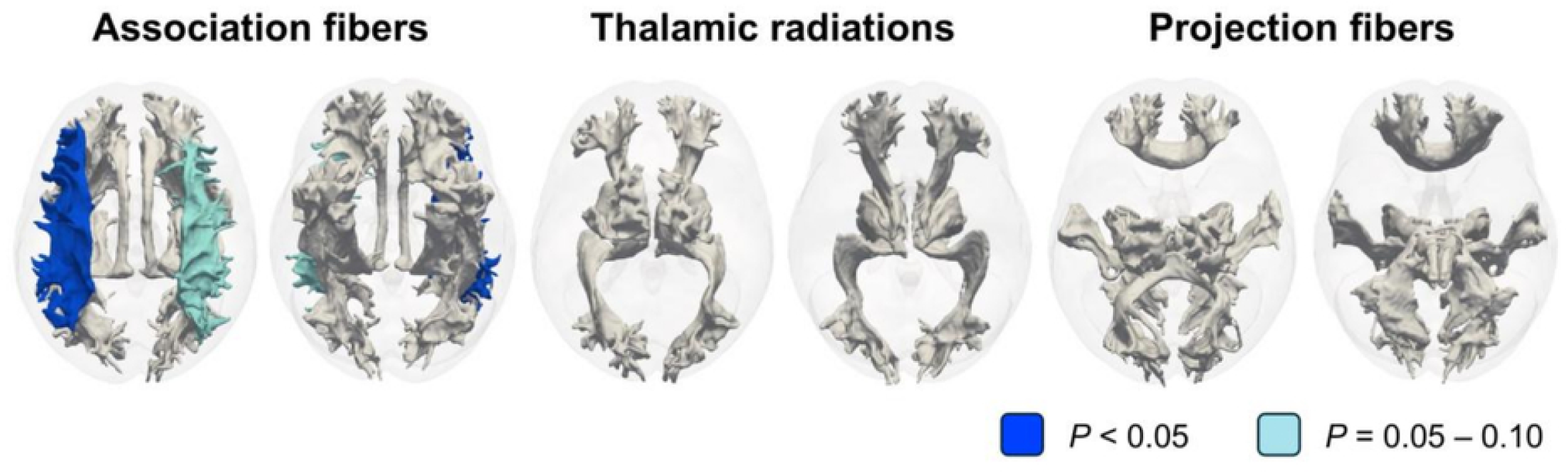
Illustrative representation of inosine triphosphate pyrophosphatase (ITPA) predicted loss-of-function burden for fractional anisotropy in each 27 white matter tract by GeneBass analysis. Probabilistic tractographies colored by Sequence Kernel Association Test–Optimal (SKAT-O) P values after Bonferroni correction for 27 multiple comparisons were shown in superior and inferior views (n=16,496). The corrected *P* value of 0.05 corresponds to a raw P value of 0.00185. White-colored tracts indicate adjusted tracts corrected *P*>0.10.

### Higher expression of ITPA in the cortex of CHF rat

We performed the Western blot analysis of rat brain tissue to investigate the association between CHF and ITPA expression. In the pericallosal regions, where FA and MBP were measured in the imaging and immunohistochemical analyses, ITPA expression was higher in the CHF group compared to the sham group (P=0.009, **Figure 5A**). The immunohistochemical analysis showed that ITPA immunoreactivity was localized around DAPI-positive nuclei and co-localised with neuronal cells (**Figure 5B**), while glial cell markers assessed in the examined sections did not co-localize with ITPA immunoreactivity in the CHF rat brain (**Figure 5C and D**).

**Figure 5.**
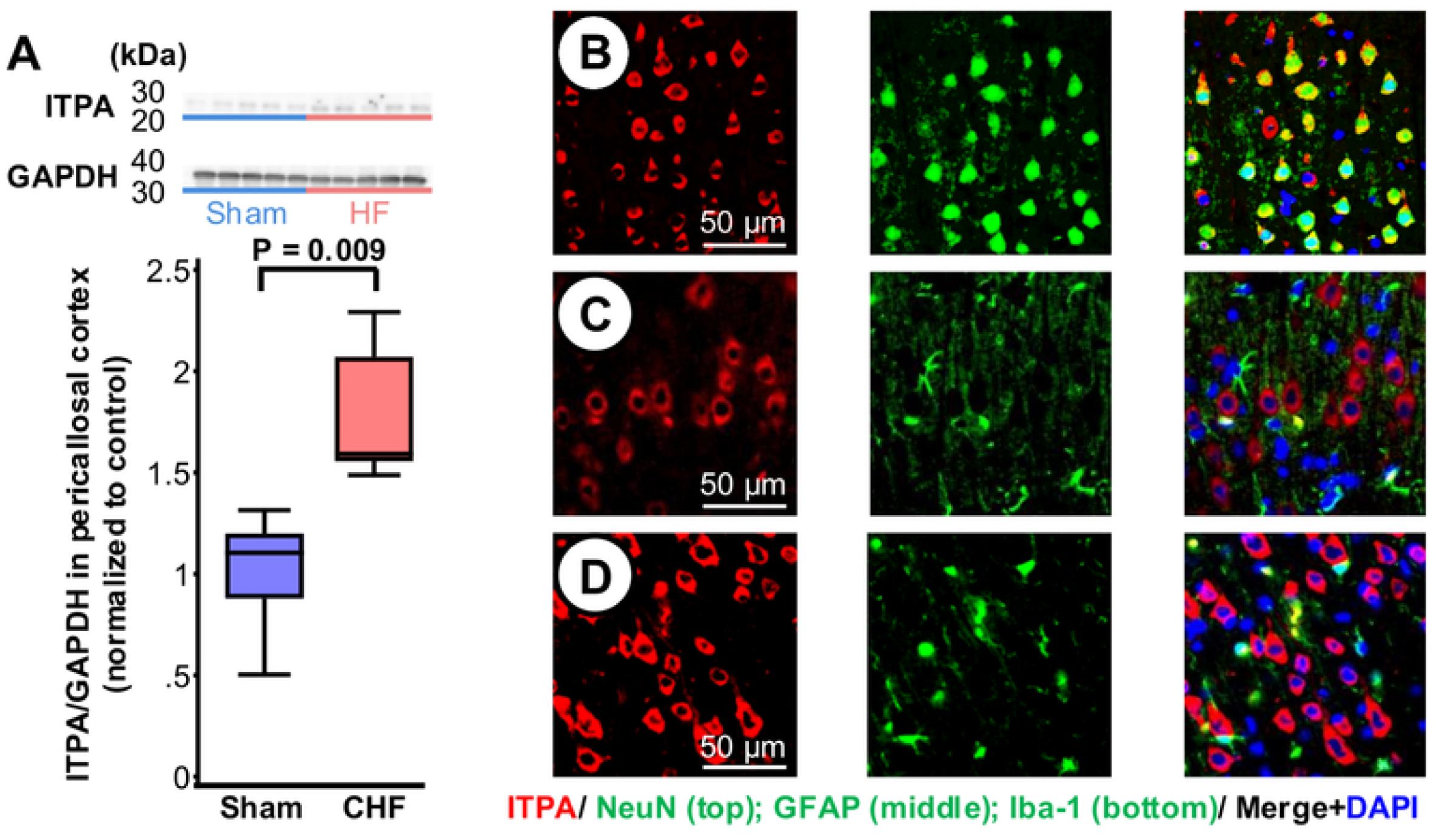
Inosine triphosphate pyrophosphatase (ITPA) expression in the rat cortex. Western blots and quantification of ITPA protein expression relative to GAPDH in the cortical region adjacent to the corpus callosum of sham and chronic heart failure rats (n=5, each). Statistical analysis was performed using Mann–Whitney U test (A). Representative immunofluorescence images of ITPA (red), neuronal nuclei (NeuN; green), glial fibrillary acidic protein (GFAP; green), Ionized calcium-binding adapter molecule 1 (Iba-1; green) and 4‘,6-diamidino-2-phenylindole (DAPI; blue) in the rat cortex. Scale bar; 50μm (B, C, D).

### Axonal injury induced by ITPA

To further investigate the effects of ITPA on WMH, we performed additional validation analyses using a cell culture. We performed morphometric analysis of cultured mouse neurons to investigate the effects of recombinant ITPA on axons, a major structural component of cerebral white matter. The immunocytochemical analysis showed that SMI31-positive axonal processes were spatially distinct from MAP2-positive somatodendritic processes (**Figure 6A and 6B**). The normalized total length of SMI31-positive axons per field was significantly decreased in the recombinant ITPA-treated group compared to the vehicle-treated group (0.648±0.217 vs. 1.000±0.379, P=0.023, **Figure 6C**).

**Figure 6.**
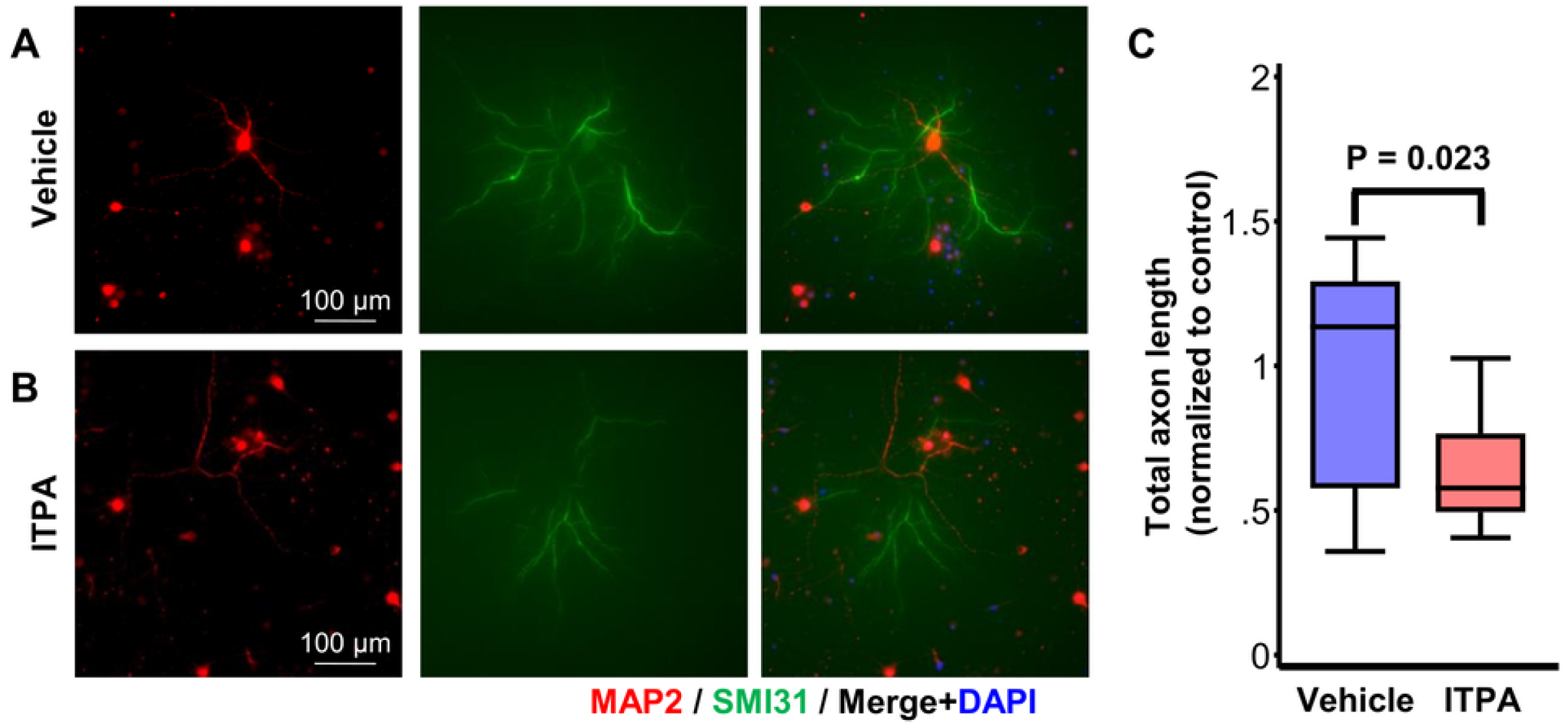
Axon damage following inosine triphosphate pyrophosphatase (ITPA) in cultured cortical neuron. Representative immunofluorescence images of microtubule-associated protein 2 (MAP2; red), phosphorylated neurofilament heavy chain (SMI31, green) and 4‘,6-diamidino-2-phenylindole (DAPI; blue) with vehicle or recombinant ITPA. Scale bar; 100μm (A, B). Quantification of the total axon length of E18 cortex neurons grown in cultures treated with vehicle or 500ng/mL recombinant ITPA normalized to the average length of vehicle (n=10, each). Statistical analysis was performed using Student’s t-test (C).

## Discussion

The key findings of this study are as follows: (1) patients with Stage C CHF showed the association with greater WMH volume than those with Stage B CHF, particularly in the anterior periventricular white matter. The CHF-associated white matter abnormalities were also observed in the rodent CHF model; (2) proteomic and MR analyses provided supportive evidence for a potential role of ITPA linking CHF and WMH progression; (3) ITPA protein expression of the pericallosal cortex was higher in CHF compared to sham rats; (4) ITPA induced axonal injury in vitro, which was consistent with the ITPA-predicted loss-of function burden for white matter FA. Collectively, these results indicate that ITPA, a circulating humoral factor, may be associated with the WMH progression in patients with CHF.

### WMH and white matter abnormalities in CHF

Various structural brain changes have been reported in patients with CHF. Gray matter loss was associated with poorer cognitive function.^37^ A greater total WMH volume is characterized by more advanced NYHA functional class and reduced LVEF,^16^ and an association between reduced cardiac output and increased WMH has also been reported.^15^ Previous DTI study have demonstrated microstructural white matter abnormalities, including alterations in axonal integrity and myelin structure, in patients with heart failure.^38^ However, the relationship between cardiac function and WMH has not been consistent across studies. Longitudinal brain MRI acquisitions in this study showed that the WMH change was greater in Stage C compared to Stage B groups. Moreover, the corresponding animal experiments showed lower FA and MBP immunodensity in the anterior pericallosal region of CHF rats, closely mirroring the regional distribution observed in the CHF patient cohort. Clinical and experimental observations have been reported in hippocampal injury of CHF in human.^5,34^ Although the mechanism is unclear, a previous study has reported that WMH in the anterior periventricular region was frequently associated with blood pressure traits.^39^ The consistent clinical and experimental animal findings of this study supported the WMH progession and underlying abnormalities in white matter microstructure in CHF.

Cerebral atherosclerosis and resultant hypoperfusion have been reported to contribute to the development of WMH in previous studies,^11,12^ suggesting low cardiac output may also be involved in this process. However, WMH progression was observed without a difference in cardiac and systolic volume indices between Stage B and Stage C groups in this study. In a German cohort of patients with heart failure with preserved ejection fraction, LVEF was not significantly associated with the Fazekas score, a semiquantitative measure of WMH burden.^17^ No significant association was observed between cardiac indices and WMH burden in the middle-aged general population.^40^ The present study does not exclude a contribution of cerebral hypoperfusion, but suggest that conventional cardiac hemodynamic measures alone may not fully explain the observed WMH progression. Other factors associated with CHF, including neurohumoral or systemic processes, may therefore warrant further investigation as potential contributors to white matter abnormalities.

### ITPA as a candidate factor in WMH progression in CHF

The mechanisms linking CHF to progressive white matter abnormalities remain incompletely understood. Focusing on the possibility that systemic molecular signals influence cerebral small vessel disease, this study explored ITPA as a possible candidate protein in the CHF-associated WMH progression through a combination of proteomic screening and MR analyses. ITPA is a house-cleaning enzyme that prevents the incorporation of noncanonical purine nucleotides into DNA and RNA by catalyzing the hydrolysis of (deoxy) nucleoside triphosphates into the corresponding nucleoside monophosphate with the concomitant release of pyrophosphate.^41^ Previous northern blot analysis demonstrated that ITPA mRNA is widely expressed across systemic human tissues including the brain, which is consistent with its role as a broadly expressed nucleotide-pool sanitizing enzyme.^42^ In humans, biallelic loss-of-function variants in ITPA have been shown to cause early infantile epileptic encephalopathy.^43^ Indeed, ITPA-knockout leads to the accumulation of inosine nucleotides in both the nucleotide and RNA pool, thus ITPA-knockout mice exhibit growth retardation and die within several weeks after birth.^44^ Consequently, studies evaluating the physiological and pathological roles of ITPA in mature animals using knockout mouse models remain limited.

This study showed that the cerebral expression of ITPA, which was localized predominantly in the cytoplasm of neurons, was higher in a rat model of CHF. Several previous studies suggest that CHF induces extensive transcriptional and epigenetic remodeling in the brain.^45^ The higher cerebral expression of ITPA observed in this study may represent a compensatory response to CHF–induced cellular stress and disturbances in the nucleotide pool. In CHF patients, the expression of numerous miRNAs has been altered, potentially modifying the relationship between target mRNA abundance and protein expression through changes in miRNA stability and translational efficiency.^46^ Therefore, alterations in miRNAs potentially involved in the regulation of ITPA expression may also have contributed to the higher cerebral expression of ITPA. Further studies should be warranted to elucidate the mechanisms underlying ITPA expression in CHF.

The results of this study suggested that ITPA, which normally exerts a cytoprotective effect by eliminating noncanonical nucleotides, contributes to neuronal injury. Although ITPA is physiologically essential, it may have detrimental effects when present in excess. In addition, whereas complete or severe loss of ITPA function causes harmful damage, a modest ITPA reduction during adulthood has been repored to confer protective outcomes under certain conditions. Patients carrying genetic variants associated with reduced ITPA activity exhibit a delayed onset of ribavirin-induced hemolytic anemia during treatment for chronic hepatitis C.^47^ The protective effect of ITPA polymorphisms is likely mediated by modulation of intracellular nucleotide pools.^48^ Furthermore, both reduced and excessive expression of certain proteins, such as TDP-43, have been shown to impair neuronal function and viability.^49^ Considering these insights of protein activity, it is conceivable that neuronal ITPA activity has an optimal phisiological range and both insufficient and excessive ITPA levels may disrupt nucleotide-pool homeostasis required for axonal maintenance, thereby potentially contributing to WMH progression. Further experimental studies are needed to determine whether such a non-linear or converse relationship underlies the association between ITPA and white matter abnormalities observed in the present study.

### Clinical implication and Future Directions

This study partially revealed a previously underexplored mechanism contributing to the development of cerebral small vessel disease in CHF. By focusing on humoral factors as a pathway that cannot be fully explained by conventional hemodynamics alone, we identified ITPA as a candidate molecular mediator linking CHF to the WMH progression. ITPA may serve as a supportive biomarker for stratifying the risk of WMH progression in patients with CHF. Furthermore, modulation of pathologically increased ITPA activity or the resulting downstream disturbances in nucleotide-pool homeostasis may provide a novel therapeutic strategy for attenuating CHF–associated white matter injury and potentially ameliorating neuropsychiatric complications of CHF, including cognitive impairment and depression. Because research using human brain tissue is limited by its invasiveness, future studies should focus on evaluation with blood and other accessible tissue samples.

### Limitations

This study has several limitations. First, the B-HeFT2 was a single-center cohort comprising only patients with CHF, which may limit the generalizability of our findings. Further validation in larger, multicenter cohorts that include appropriate control groups is warranted. Second, the assessment of ITPA was limited to aptamer-based proteomic analysis of human plasma and analysis of brain tissue from the animal model of CHF, thus circulating ITPA and cerebral ITPA were not linked directly. Future studies should accurately quantify concentrations of circulating ITPA. Third, the statistical significance of proteomic and MR analyses was exploratory although independent evidence from animal model and community-based cohort was consistent. Fourth, the development of cerebral white matter differs between humans and rodents. White matter occupies a greater proportion of cerebral volume in humans than in small rodents, thus the scaling relationships between white matter volume and cortical connectivity differ across mammalian species.^50^

### Conclusions

Symptomatic CHF was associated with accelerated WMH progression, including white matter abnormalities such as lower FA. Complementary experimental and exploratory genetic findings identify ITPA as a candidate protein for the biological pathways linking CHF-associated white matter abnormalities. The results provide a basis for further investigation of ITPA as a novel therapeutic target for brain health.

## Acknowledgments

The authors would like to thank Ms. Watanabe and Mr. Kondo for their technical support in animal experiment. This study was supported by Support system for young researchers to use research equipment, instruments and devices in Tohoku University.

## Data Availability

The UK Biobank-derived exome association summary statistics used in this study are publicly available through the GeneBass database (https://app.genebass.org/). The other data underlying this article are available from the corresponding author upon reasonable request.

## References

1. Writing Committee Members, ACC/AHA Joint Committee Members. 2022 AHA/ACC/HFSA Guideline for the Management of Heart Failure. J Card Fail. 2022;28:e1–e167.

2. Qiu C, Winblad B, Marengoni A, Klarin I, Fastbom J, Fratiglioni L. Heart Failure and Risk of Dementia and Alzheimer Disease: A Population-Based Cohort Study. Arch Intern Med. 2006;166:1003–1008.

3. Rutledge T, Reis VA, Linke SE, Greenberg BH, Mills PJ. Depression in Heart Failure. JACC. 2006;48:1527–1537.

4. Suzuki H, Sumiyoshi A, Matsumoto Y, Duffy BA, Yoshikawa T, Lythgoe MF, Yanai K, Taki Y, Kawashima R, Shimokawa H. Structural abnormality of the hippocampus associated with depressive symptoms in heart failure rats. NeuroImage. 2015;105:84–92.

5. Suzuki H, Matsumoto Y, Ota H, Sugimura K, Takahashi J, Ito K, Miyata S, Furukawa K, Arai H, Fukumoto Y, et al. Hippocampal Blood Flow Abnormality Associated With Depressive Symptoms and Cognitive Impairment in Patients With Chronic Heart Failure. Circ J. 2016;80:1773–1780.

6. Suzuki H, Matsumoto Y, Ota H, Sugimura K, Takahashi J, Ito K, Miyata S, Fukumoto Y, Taki Y, Shimokawa H. Structural brain abnormalities and cardiac dysfunction in patients with chronic heart failure. European Journal of Heart Failure. 2018;20:936–938.

7. Debette S, Markus HS. The clinical importance of white matter hyperintensities on brain magnetic resonance imaging: systematic review and meta-analysis. BMJ. 2010;341:c3666–c3666.

8. Debette S, Schilling S, Duperron M-G, Larsson SC, Markus HS. Clinical Significance of Magnetic Resonance Imaging Markers of Vascular Brain Injury. JAMA Neurol. 2019;76:81–94.

9. Wardlaw JM, Smith C, Dichgans M. Small vessel disease: mechanisms and clinical implications. The Lancet Neurology. 2019;18:684–696.

10. Hase Y, Horsburgh K, Ihara M, Kalaria RN. White matter degeneration in vascular and other ageing-related dementias. Journal of Neurochemistry. 2018;144:617–633.

11. Joutel A, Chabriat H. Pathogenesis of white matter changes in cerebral small vessel diseases: beyond vessel-intrinsic mechanisms. Clinical Science. 2017;131:635–651.

12. Liu Y, Dong Y-H, Lyu P-Y, Chen W-H, Li R. Hypertension-Induced Cerebral Small Vessel Disease Leading to Cognitive Impairment. Chinese Medical Journal. 2018;131:615–619.

13. Sargurupremraj M, Suzuki H, Jian X, Sarnowski C, Evans TE, Bis JC, Eiriksdottir G, Sakaue S, Terzikhan N, Habes M, et al. Cerebral small vessel disease genomics and its implications across the lifespan. Nat Commun. 2020;11:6285.

14. Solé-Guardia G, Custers E, De Lange A, Clijncke E, Geenen B, Gutierrez J, Küsters B, Claassen JAHR, De Leeuw F-E, Wiesmann M, et al. Association between hypertension and neurovascular inflammation in both normal-appearing white matter and white matter hyperintensities. acta neuropathol commun. 2023;11:2.

15. Jefferson AL, Tate DF, Poppas A, Brickman AM, Paul RH, Gunstad J, Cohen RA. Lower Cardiac Output Is Associated with Greater White Matter Hyperintensities in Older Adults with Cardiovascular Disease. J Am Geriatr Soc. 2007;55:1044–1048.

16. Vogels RLC, Van Der Flier WM, Van Harten B, Gouw AA, Scheltens P, Schroeder-Tanka JM, Weinstein HC. Brain Magnetic Resonance Imaging Abnormalities in Patients with Heart Failure. European Journal of Heart Failure. 2007;9:1003–1009.

17. Stegmann T, Chu ML, Witte VA, Villringer A, Kumral D, Riedel-Heller SG, Roehr S, Hagendorff A, Laufs U, Loeffler M, et al. Heart Failure is Independently Associated with White Matter Lesions: Insights from the Population-Based LIFE-Adult Study. ESC Heart Failure. 2021;8:697–704.

18. Maillard P, Fletcher E, Harvey D, Carmichael O, Reed B, Mungas D, DeCarli C. White Matter Hyperintensity penumbra. Stroke. 2011;42:1917–1922.

19. 2005 WRITING COMMITTEE MEMBERS, Hunt SA, Abraham WT, Chin MH, Feldman AM, Francis GS, Ganiats TG, Jessup M, Konstam MA, Mancini DM, et al. 2009 Focused Update Incorporated Into the ACC/AHA 2005 Guidelines for the Diagnosis and Management of Heart Failure in Adults. Circulation. 2009;119:e391–e479.

20. Shah AM, Myhre PL, Arthur V, Dorbala P, Rasheed H, Buckley LF, Claggett B, Liu G, Ma J, Nguyen NQ, et al. Large scale plasma proteomics identifies novel proteins and protein networks associated with heart failure development. Nat Commun. 2024;15:528.

21. Gold L, Ayers D, Bertino J, Bock C, Bock A, Brody EN, Carter J, Dalby AB, Eaton BE, Fitzwater T, et al. Aptamer-Based Multiplexed Proteomic Technology for Biomarker Discovery. PLoS One. 2010;5:e15004.

22. Kurki MI, Karjalainen J, Palta P, Sipilä TP, Kristiansson K, Donner KM, Reeve MP, Laivuori H, Aavikko M, Kaunisto MA, et al. FinnGen provides genetic insights from a well-phenotyped isolated population. Nature. 2023;613:508–518.

23. Ferkingstad E, Sulem P, Atlason BA, Sveinbjornsson G, Magnusson MI, Styrmisdottir EL, Gunnarsdottir K, Helgason A, Oddsson A, Halldorsson BV, et al. Large-scale integration of the plasma proteome with genetics and disease. Nat Genet. 2021;53:1712–1721.

24. Mustapha M, Nassir CMNCM, Aminuddin N, Safri AA, Ghazali MM. Cerebral Small Vessel Disease (CSVD) – Lessons From the Animal Models. Front Physiol. 2019;10:1317.

25. Antonsen BT, Jiang Y, Veraart J, Qu H, Nguyen HP, Sijbers J, von Hörsten S, Johnson GA, Leergaard TB. Altered diffusion tensor imaging measurements in aged transgenic Huntington disease rats. Brain Struct Funct. 2013;218:767–778.

26. Reep RL. Cortical Layer VII and Persistent Subplate Cells in Mammalian Brains. Brain Behav Evol. 2000;56:212–234.

27. Watakabe A, Hirokawa J. Cortical networks of the mouse brain elaborate within the gray matter. Brain Struct Funct. 2018;223:3633–3652.

28. Yushkevich PA, Piven J, Hazlett HC, Smith RG, Ho S, Gee JC, Gerig G. User-guided 3D active contour segmentation of anatomical structures: significantly improved efficiency and reliability. Neuroimage. 2006;31:1116–1128.

29. Karczewski KJ, Solomonson M, Chao KR, Goodrich JK, Tiao G, Lu W, Riley-Gillis BM, Tsai EA, Kim HI, Zheng X, et al. Systematic single-variant and gene-based association testing of thousands of phenotypes in 394,841 UK Biobank exomes. Cell Genomics. 2022;2:100168.

30. Suzuki H, Gao H, Bai W, Evangelou E, Glocker B, O’Regan DP, Elliott P, Matthews PM. Abnormal brain white matter microstructure is associated with both pre-hypertension and hypertension. PLoS ONE. 2017;12:e0187600.

31. Lee S, Emond MJ, Bamshad MJ, Barnes KC, Rieder MJ, Nickerson DA, Christiani DC, Wurfel MM, Lin X. Optimal Unified Approach for Rare-Variant Association Testing with Application to Small-Sample Case-Control Whole-Exome Sequencing Studies. Am J Hum Genet. 2012;91:224–237.

32. Madan CR. Creating 3D visualizations of MRI data: A brief guide. F1000Res. 2015;4:466.

33. Chen C, Hayden KM, Kaufman JD, Espeland MA, Whitsel EA, Serre ML, Vizuete W, Orchard TS, Wang X, Chui HC, et al. Adherence to a MIND-Like Dietary Pattern, Long-Term Exposure to Fine Particulate Matter Air Pollution, and MRI-Based Measures of Brain Volume: The Women’s Health Initiative Memory Study-MRI. Environ Health Perspect. 2021;129:127008.

34. Suzuki H, Matsumoto Y, Sugimura K, Takahashi J, Miyata S, Fukumoto Y, Taki Y, Shimokawa H. Impacts of hippocampal blood flow on changes in left ventricular wall thickness in patients with chronic heart failure. International Journal of Cardiology. 2020;310:103–107.

35. von Elm E, Altman DG, Egger M, Pocock SJ, Gøtzsche PC, Vandenbroucke JP, STROBE Initiative. The Strengthening the Reporting of Observational Studies in Epidemiology (STROBE) statement: guidelines for reporting observational studies. PLoS Med. 2007;4:e296.

36. Skrivankova VW, Richmond RC, Woolf BAR, Yarmolinsky J, Davies NM, Swanson SA, VanderWeele TJ, Higgins JPT, Timpson NJ, Dimou N, et al. Strengthening the Reporting of Observational Studies in Epidemiology Using Mendelian Randomization: The STROBE-MR Statement. JAMA. 2021;326:1614–1621.

37. Almeida OP, Garrido GJ, Beer C, Lautenschlager NT, Arnolda L, Flicker L. Cognitive and brain changes associated with ischaemic heart disease and heart failure. Eur Heart J. 2012;33:1769–1776.

38. Kumar R, Woo MA, Macey PM, Fonarow GC, Hamilton MA, Harper RM. Brain Axonal and Myelin Evaluation in Heart Failure. J Neurol Sci. 2011;307:106–113.

39. Beyer F, Tsuchida A, Soumaré A, Rajula HSR, Mishra A, Crivello F, Proust-Lima C, Loeffler M, Tzourio C, Amouyel P, et al. White matter hyperintensity spatial patterns: Risk factors and clinical correlates. Alzheimer’s & Dementia. 2025;21:e70053.

40. van Hout MJP, Dekkers IA, Westenberg JJM, Schalij MJ, Scholte AJHA, Lamb HJ. Associations between left ventricular function, vascular function and measures of cerebral small vessel disease: a cross-sectional magnetic resonance imaging study of the UK Biobank. Eur Radiol. 2021;31:5068–5076.

41. Zamzami MA. Inosine Triphosphate Pyrophosphatase (ITPase): Functions, Mutations, Polymorphisms and Its Impact on Cancer Therapies. Cells. 2022;11:384.

42. Lin S, McLennan AG, Ying K, Wang Z, Gu S, Jin H, Wu C, Liu W, Yuan Y, Tang R, et al. Cloning, expression, and characterization of a human inosine triphosphate pyrophosphatase encoded by the itpa gene. J Biol Chem. 2001;276:18695–18701.

43. Scala M, Wortmann SB, Kaya N, Stellingwerff MD, Pistorio A, Glamuzina E, Karnebeek CD, Skrypnyk C, Iwanicka-Pronicka K, Piekutowska-Abramczuk D, et al. Clinico-radiological features, molecular spectrum, and identification of prognostic factors in developmental and epileptic encephalopathy due to inosine triphosphate pyrophosphatase (ITPase) deficiency. Human Mutation. 2022;43:403–419.

44. Behmanesh M, Sakumi K, Abolhassani N, Toyokuni S, Oka S, Ohnishi YN, Tsuchimoto D, Nakabeppu Y. ITPase-deficient mice show growth retardation and die before weaning. Cell Death Differ. 2009;16:1315–1322.

45. Islam MR, Lbik D, Sakib MS, Maximilian Hofmann R, Berulava T, Jiménez Mausbach M, Cha J, Goldberg M, Vakhtang E, Schiffmann C, et al. Epigenetic gene expression links heart failure to memory impairment. EMBO Mol Med. 2021;13:EMMM201911900.

46. Peterlin A, Počivavšek K, Petrovič D, Peterlin B. The Role of microRNAs in Heart Failure: A Systematic Review. Front Cardiovasc Med. 2020;7:161.

47. Thompson Alexander J, Fellay J, Patel K, Tillmann HL, Naggie S, Ge D, Urban TJ, Shianna KV, Muir AJ, Fried MW, et al. Variants in the ITPA Gene Protect Against Ribavirin-Induced Hemolytic Anemia and Decrease the Need for Ribavirin Dose Reduction. Gastroenterology. 2010;139:1181–1189.

48. Hitomi Y, Cirulli ET, Fellay J, McHutchison JG, Thompson AJ, Gumbs CE, Shianna KV, Urban TJ, Goldstein DB. Inosine Triphosphate Protects Against Ribavirin-Induced Adenosine Triphosphate Loss by Adenylosuccinate Synthase Function. Gastroenterology. 2011;140:1314–1321.

49. Diaper DC, Adachi Y, Sutcliffe B, Humphrey DM, Elliott CJH, Stepto A, Ludlow ZN, Vanden Broeck L, Callaerts P, Dermaut B, et al. Loss and gain of Drosophila TDP-43 impair synaptic efficacy and motor control leading to age-related neurodegeneration by loss-of-function phenotypes. Hum Mol Genet. 2013;22:1539–1557.

50. Ventura-Antunes L, Mota B, Herculano-Houzel S. Different scaling of white matter volume, cortical connectivity, and gyrification across rodent and primate brains. Front. Neuroanat. [Internet]. 2013 [cited 2026 July 15];7. Available from: http://journal.frontiersin.org/article/10.3389/fnana.2013.00003/abstract

